# Inferring B cell specificity for vaccines using a mixture model

**DOI:** 10.1101/464792

**Authors:** Anna Fowler, Jacob D. Galson, Johannes Trück, Dominic F. Kelly, Gerton Lunter

## Abstract

Vaccines have greatly reduced the burden of infectious disease, ranking in their impact on global health second only after clean water. Most vaccines confer protection by the production of antibodies with binding affinity for the antigen, which is the main effector function of B cells. This results in short term changes in the B Cell receptor (BCR) repertoire when an immune response is launched, and long term changes when immunity is conferred. Analysis of antibodies in the serum is usually used to evaluate vaccine response, however this is limited and therefore the investigation of the BCR repertoire provides far more detail for the analysis of vaccine response. Here, we introduce a novel Bayesian model to describe the observed distribution of BCR sequences and the pattern of sharing across time and between individuals, with the goal to identify vaccine-specific BCR sequences. We use data from two studies to assess the model and estimate that we can identify vaccine-specific sequences with 69% sensitivity.

## Introduction

The array of potential foreign antigens that the human immune system must provide protection against is vast, and an individual’s B Cell receptor (BCR) repertoire is correspondingly huge; it is estimated that a human adult has over 10^13^ theoretically possible BCRs [1], of which as many as 10^11^ may be realized [2]. This diversity is primarily generated through recombination, junctional diversity, and somatic mutation of the V, D and J segments of the immunoglobulin heavy chain genes (IgH) [2], combined with selection to avoid self-reactivity and to increase antigen specificity. The BCR repertoire of a healthy individual is constantly evolving, through the generation of novel naive B cells, and by the maturation and activation of B cells stimulated by ongoing challenges of pathogens and other antigens. As a result, an individual’s BCR repertoire is unique and dynamic, and is influenced by age, health and infection history as well as genetic background [3].

Upon stimulation, B cells undergo a process of proliferation and hyper-mutation, resulting in the selection of clones with improved antigen binding and the ability to mount an effective immune response. The process of hyper-mutation targets specific regions, and subsequent selection provides a further focusing of sequence changes. The short genomic region in which most of these changes occur, and which is thought to play a key role in determining antigen binding specificity, is termed the Complementarity Determining Region 3 (CDR3) [4, 5]. Next generation sequencing (NGS) makes it possible to capture the CDR3 across a large sample of cells, providing a sparse but high-resolution snapshot of the BCR repertoire, and forming a starting point to study immune response and B-cell-mediated disease [6].

Vaccination provides a controlled and easily administered stimulus that can be used to study this complex system [7]. An increase in clonality has been observed in the post-vaccination BCR repertoire, which has been related to the proliferation of B cells and the production of active plasma cells [8–14]. An increase in the sequences shared between individuals, referred to as the public repertoire or stereotyped BCRs, has also been observed, and there is mounting evidence that this public repertoire is at least partly due to convergent evolution in different individuals responding to the same stimulus [10, 14–18].

These observations suggest that by identifying similarities between the BCR repertoires of a group of individuals that have received a vaccine stimulus, it may be possible to identify B cells specific to the vaccine. However, while the most conspicuous of these signals could be shown to be likely due to a convergent response to the same antigen in multiple individuals [19], it is much harder to link more subtle signals to vaccine response using ad-hoc classification methods. To address this, we here develop a statistical model for the abundance of BCRs over time in multiple individuals, which integrates the signals of increased expression, clonality, and sharing across individuals. We use this model to classify BCRs into three classes depending on the inferred states of their B cell hosts, namely non-responders (background, bg), those responding to a stimulus other than the vaccine (non-specific, ns), and those responding to the vaccine (vaccine-specific, vs).

Here we show that the sequences classified as vaccine-specific by our model have distinct time profiles and patterns of sharing between individuals, and are enriched for sequences derived from B cells that were experimentally enriched for vaccine specificity. Moreover, we show that sequences identified as vaccine-specific cluster in large groups of high sequence similarity, a pattern that is not seen in otherwise similar sets of sequences.

## Methods

### BCR Repertoire Vaccine Study Data Sets

We use two publicly available data sets, one from a study involving a hepatitis-B vaccine [20] and one from astudy on an influenza vaccine [10]. We describe these two data sets below. Both data sets capture the somatically rearranged VDJ region in B cells, in particular the highly variable CDR3 region on which we will focus.

### Hepatitis B

In the study by Galson and colleagues [20], 5 subjects were given a booster vaccine against hepatitis B (HepB) following an earlier primary course of HepB vaccination. Samples were taken on days 0, 7, 14, 21 and 28 relative to the day of vaccination. Total B cells were sorted and sequenced in all samples. We refer to this data set as the *hepatitis B data set*.

In addition, cells were sorted for HepB surface antigen specificity at the same time points post-vaccination, and then sequenced. These cells are enriched with those we are seeking to identify using our modelling approach, and provides a way to validate our method. We refer to these data as the *HBsAG+ data set*. Both data sets are publicly available on the Short Read Archive (accession PRJNA308641).

Sequences were generated on the Illumina platform using an RNA sequencing protocol, and the nucleotide sequences analysed. Targeting RNA means that highly abundant sequences may derive either from multiple B cells from a clonal subpopulation, or from one or a small number of with high IgH gene expression, such as plasma cells that are actively secreting antibodies. Although we cannot distinguish between these two possibilities, both classes of cells are likely signifiers of immune response, and are therefore of interest.

### Influenza

We also analyze data from subjects that were vaccinated against influenza in a study by Jackson and colleagues [10]. Samples were taken on days 0, 7 and 21 relative to vaccination. We analyzed a subset of 7 subjects that were deemed to be “seroconverters” based on vaccine-specific ELISA assays. This will be referred to as the *influenza data set*.

In addition, the authors also collected plasmablasts on day 7 in 5 of the subjects. These are also likely to be enriched for B cells responding to the vaccine and therefore provide an additional source of evaluation for our method. The sequences derived from these cells are referred to as the *plasmablast data set*. All data is publicly available on dbGaP (accession phs000760.v.1.p1).

The Roche 454 platform was used to perform DNA sequencing of the somatically recombined IgH locus, using primers for the relatively conserved FR2 IgH V gene segment, and a conserved J gene segment, and we analyse the amino acid sequences. Targeting DNA ensures that sequences with high abundance are representative of clonally expanded B cells, rather than of cells exhibiting high mRNA expression. However, active plasma cells with high secretion rate would still be counted individually.

### Clustering

We combined sequences into *clusters*, in order mainly to correct for read errors, as well as to group together some highly similar sequences that likely target the same epitope. This removes some noise associated with read error and strengthens signals by treating multiple sequences all of which target the same epitope as a single cluster, whilst also reducing the computational burden. Each cluster consists of a single identifying CDR3 sequence, the *cluster centre*, and its’ set of neighbouring CDR3 sequences; for two sequences to be considered neighbours, they must be of the same length and be highly similar, which we define as greater than 85% similarity for nucleotide sequences as in the hepatitis B data set, or 90% similarity for amino acid sequences as in the influenza data set. The clustering was performed in a greedy manner, by iteratively identifying a cluster centre as the sequence with the greatest number of neighbours from among all unclustered sequences, and assigning it and its unclustered neighbours to a new cluster. This is a computationally efficient approach to clustering which allows us to cluster very large data sets. However, the model presented here is not dependent on the clustering algorithm used, and any alternative clustering method could also be used as input.

Within each data set, we clustered sequences from all samples and time points together, but kept track of sample- and time-specific counts to enable the analysis of time dynamics and between-individual sharing. We now consider each cluster to be representative of the sequence *i* at its centre, and make no distinction between clusters and the individual sequences which form the cluster centres. In addition we shall use *i* to refer to the B cell(s) or B cell clone that the cluster represents. We define the *sequence abundance*, denoted by *x_ist_*, as the number of sequences assigned to the cluster represented by sequence *i* for a participant *s* at time point *t*, and the total sequence abundance as the total number of sequences assigned to the cluster across all samples, Σ*_st_x_ist_*.

### Model

We introduce a hierarchical Bayesian model to describe the abundance of BCR sequences (or alternatively, CDR3 sequences) across individuals inoculated with the same vaccine, and across multiple time points. The data are abundances, *x_ist_*, as introduced above. The goal of modeling these data is to identify vaccine-specific sequences from among a large number of non-vaccine-specific sequences, while accounting for the sparse sampling of the sequences and for the highly stochastic nature of the biological process that generates them.

One identifying feature of vaccine-specific sequences that we want to model is their abundance profile. We expect to observe no vaccine-specific sequences pre-vaccination (or very few, in the case of a primer-boost design such as for the HepB data set), while post-vaccination we expect to observe high abundances due to clonal expansion of stimulated B cells, the presence of plasma cells with high transcription activity, or both. A second feature that helps to characterise vaccine-specific sequences is their tendency to be shared across individuals, due to convergent evolution.

To describe the model we introduce some notation. As above let *i* denote a sequence, and denote by Ω the space of all sequences. We partition this set as Ω = Ω_bg_ ∪ Ω_vs_ ∪ Ω_ns_, where the disjunct subsets represent background sequences not responding to any stimulus; vaccine-specific sequences responding to the vaccine stimulus; and sequences responding to a non-specific stimulus other than the vaccine respectively. These subsets (and their sizes) are unknown, and the classification of a particular sequence *i* is given by a discrete random variable γ_*i*_ ∈ {`bg, vs, ns`}, so that *i* ∈ Ω_*γi*_.

Next, the presence of a particular B cell (or cluster of B cells) *i* in a participant *s* is encoded by a second discrete random variable *z_is_*, which takes on the value 0 when *i* is absent from the BCR repertoire of individual *s* at any time point, and 1 when *i* is present in the individual (though not necessarily present in any sample taken from this individual). The variable *z* aims to account for the sparsity resulting from the diversity of BCR repertoires from different individuals. The distribution of *z_is_* is dependent on *_i_*, to allow modeling the increased probability that vaccine-specific sequences are shared between individuals.

The actual abundances *x_ist_* of sequence *i* in individual *s* at a time point *t* are assumed to be independent conditional on γ*_i_* and *z_is_*, and are modeled by a mixture of three distributions representing three outcomes, modeled by a third discrete random variable *e_ist_* whose distribution depends on γ*_i_*, *z_is_* and *t*. First, the relevant B cell or cells may be absent from individual *s* (if *z_is_* = 0) or may have escaped sampling. In this case *x_ist_* is distributed as a point mass at 0. Second, if B cells have been sampled, they may be neither clonal nor plasma B cells, and would therefore contribute a small number of sequences to the data set. In this case *x_ist_* is modeled as a negative Binomial distribution. The remaining case is that the sampled B cell or cells are either plasma cells, or cells sampled from a large clonal population (or both), in which case they are expected to contribute a large number of sequences. In this case *x_ist_* is modeled as a discretised generalized Pareto distribution [21]. This distribution of abundances is illustrated in Figure 1a. The mixture distribution of sequence counts *x_ist_* is given by *p*(*x_ist_|e_ist_, **θ***), where ***θ*** is the vector of parameters of the negative Binomial and generalized Pareto distributions.

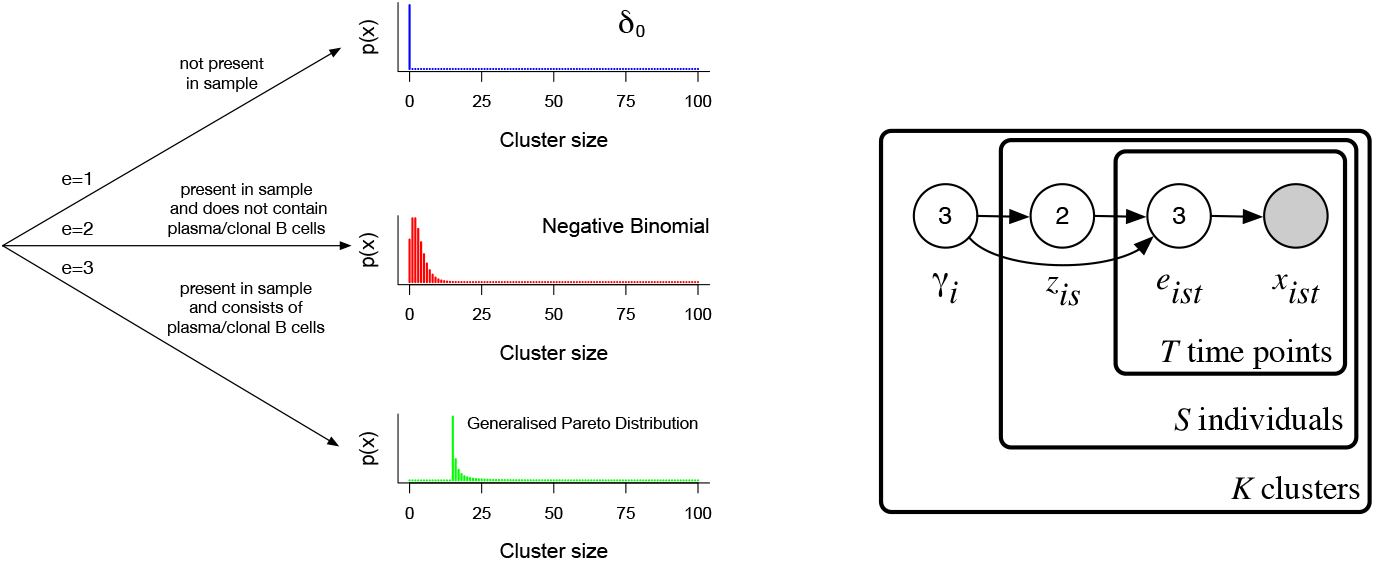
(a) Tree diagram in which each leaf represents a generative distribution for sequence abundances. The probability of following each path is dependent on the classification of the sequence and the presence of the sequence in the individual. (b) Partial graphical representation of model using plate notation. For clarity hyperparameters are not shown; Figure S1 shows a complete diagram.

The resulting joint probability for a dataset ***x***, latent variables ***e***, ***z*** and parameters **γ, *θ*** under this model is given by

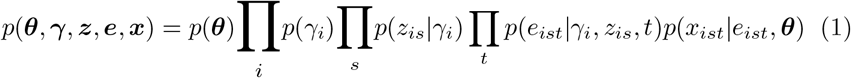

The relationship between the variables in the model is shown in Figure 1b. Non-informative priors *p*(***θ***) and *p*(γ) are placed on the parameters; this allows these parameters to be learnt from the data, and therefore allows the model to be applied to a range of data sets, for instance RNA sequencing and DNA sequencing. Full details of the model and priors are provided in the supplementary materials.

We restrict *i* to range over only those sequences which are observed at least once in the dataset, rather than the 10^13^ sequences that are theoretically possible. Therefore, for *K* sequences, we have that 1 ≤ *i* ≤ *K*. This simplifies model fitting, but will result in parameter estimates which are specific to each individual data set, and therefore affected by features such as the number of individuals. This should be kept in mind when interpreting the results.

### Inference

The model is fitted to each data set using an E-M algorithm; see the supplementary materials for details.

Restrictions on parameter values allow us to encode additional structure and to link parameters hierarchically. Firstly we assume that there is no time-dependence for the abundances of B cells classified as background or as nonspecific responders. We further assume that for the vaccine-specific cells, the pre-vaccination abundances (at *t* = 0) follow the same distribution as B cells classified as background, while post-vaccination these cells follow the same abundance distribution as B cells classified as non-specific responders. Third, we assume that the probability of a sequence being observed in a subject is the same for B cells classified as background and those classified as a non-specific response. In effect this assumes that non-specific responders are or have been responding to private stimuli, rather than for instance earlier common infections.

The uncertainty in the inferred model parameters is negligible in comparison to the biological noise because of the large amount of data. Rather than reporting this spurious precision, we report the parameter estimates without error bars, but we note that errors due to model misspecification are likely to be substantial. We report the inferred probability of a sequence belonging to each category, Γ_*class*_ for *class* ∈ {`bg, vs, ns`}. We also report, for each class, the probability that a sequence is observed given that a corresponding B cell of that class is present in an individual, *p_class_*. Finally, we report for each class the inferred probability that a sequence is being observed at high frequency, *ω_class_*.

### Sequence similarity

To compare the within-set similarity of sequences between subsets of sequences of any length, we use the Levenshtein (or “edit”) distance as implemented in [22]. Specifically, given a subset of sequences, we compute a measure of within-set similarity the mean of the Levenshtein distances between all pairs of sequences in the subset. To assess significance we use bootstrapping: we calculate the mean Levenshtein distance between a randomly selected subset of the same size, and compare the resulting null distribution of means to calculate the empirical p-value.

## Results

### Hepatitis B data set

We fitted the model to the hepatitis B data set, and obtained a good fit (see Figure S2 for the QQ plot) with key parameter estimates given in Table 1. The value of Γ_*class*_ show that most sequences are assigned to the background population, with only a small fraction responding to any stimuli. (This is also seen from the numbers shown in Table 2.) Sequences classified as vaccine specific are highly likely to be shared between multiple individuals, reflected in a high estimate of *p*_vs_, and the high estimate of *ω*_vs_ mean they are also more likely to be seen at high frequencies than those classified as background.

**Table 1:** Fitted parameters to the hepatitis B data set. Γ, the probability of a sequence belonging to each class; *p*, the probability of a sequence from each class being observed in an individual; *ω*, the probability of an observed sequence in each class being seen at high abundance.

| | $\Gamma_{class}$ | | | $p_{class}$ | | $\omega_{class}$ | |
| --- | --- | --- | --- | --- | --- | --- | --- |
| class | bg | ns | vs | bg; ns | vs | bg; $vs_{t=0}$ | ns; $vs_{t>0}$ |
|  | .992 | .005 | .003 | .216 | .970 | .006 | .277 |

**Table 2:** Number of sequences allocated to each category across all samples and the mean total sequence abundance across all samples, in the whole data set and in the subset also labelled as HBsAG+.

| Classification | All sequences |  | HBsAG+ sequences |  |
| --- | --- | --- | --- | --- |
|  | Number | Abundance (sd) | Number | Abundance (sd) |
| Background | 1,026,523 | 2.62 (31) | 60,215 | 3.45 (44) |
| Non-specific | 5,123 | 89.3 (748) | 1,500 | 147.1 (1,084) |
| Vaccine-specific | 2,976 | 10.8 (174) | 2,055 | 10.7 (190) |

For each of the three classes of sequence, the relative abundance of those sequences within individuals and the number of individuals sharing them over time are illustrated in Figure 2. The vaccine specific sequences are seen at lower frequencies at day 0 compared to subsequent time points, but still at higher frequencies than sequences classified as background. The number of individuals sharing the vaccine specific sequences increases over time up to a peak at day 14 after which sequence sharing declines again, whereas in the other classes there is no significant trend in sharing across time points, as expected.

**Figure 2:**
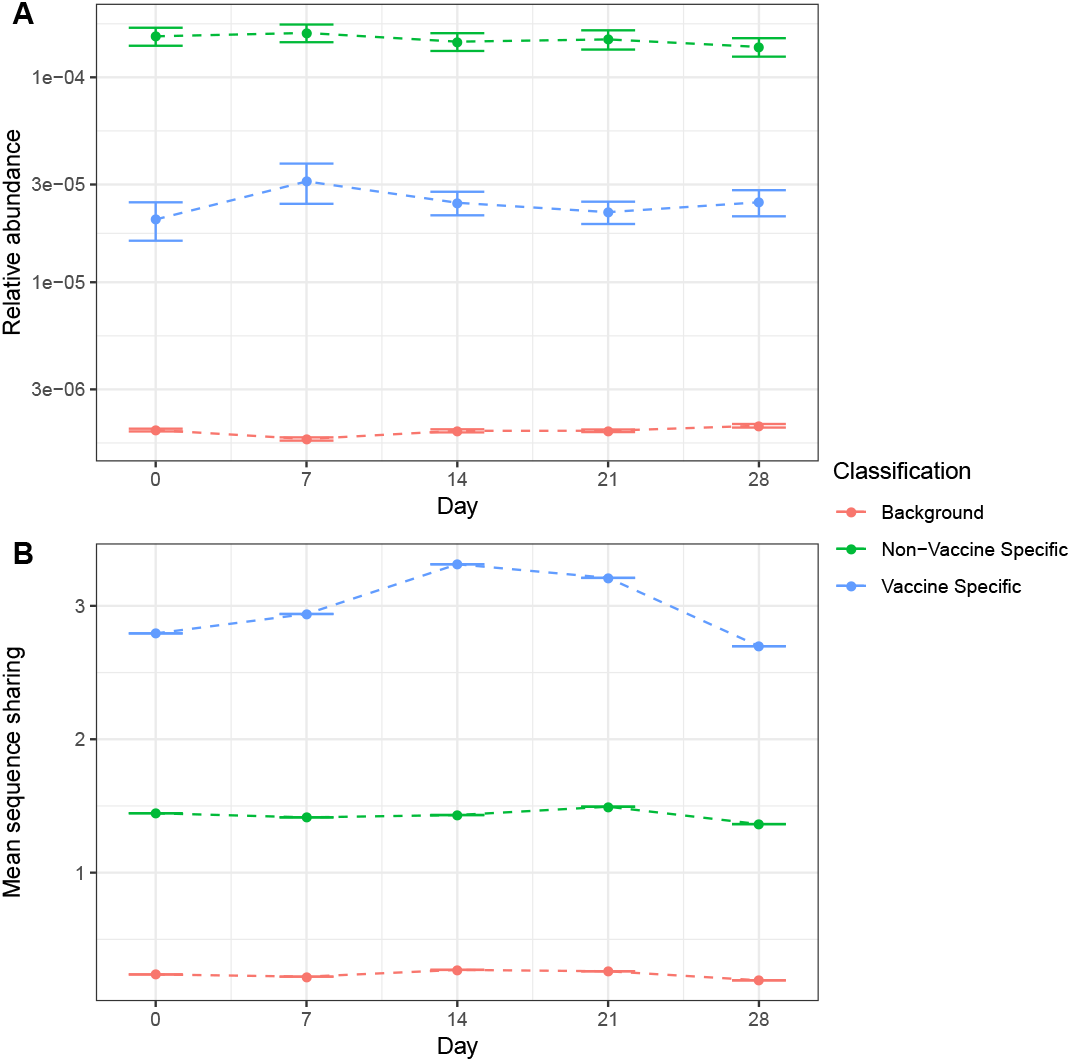
Mean sequence relative abundance at each time point in each classification (A), and the mean number of individuals sharing a sequence over time in each classification (B), for the hepatitis B data set

The total number of sequences allocated to each class and the mean total abundance of sequences from all samples within each class are shown in Table 2. Sequences are overwhelmingly classified as background, while of the remainder, similar numbers of sequences classified as non-specific responders and vaccine-specific responders. Sequences classified as background all have very low abundance, often consisting of a single sequence observed in a single individual at a single time point. Sequences classified as non-specific form the largest clusters, and are often seen at high frequency across all time points.

We next compared the hepatitis B data set with the HBsAG+ data. Sequences from the hepatitis B data set were considered present in the HBsAG+ data set if there is a sequence in the HBsAG+ data which would be assigned to the same cluster. The number of sequences from the hepatitis B data set that are present in the HBsAG+ data set, along with their abundances, are also given in Table 2. 60,215 (5.9%) of the sequences classified as background were also present in the HBsAg+ data set, however a much larger fraction (69%) of those classified as vaccine-specific were also seen in the HBsAG+ sequences. The HBsAG+ data set contains a large number of erroneously captured cells, with the specificity of staining estimated to be around 50% [20], however these erroneously captured cells are likely to be those present in high abundance in the whole repertoire (and therefore in the hepatitis B data set) due to random chance. The difference in enrichment between the background and vaccine specific categories will therefore be partly driven by the different average cluster sizes of background sequences (2.62) compared to vaccine-specific sequences (10.8). However, the fraction of non-specific responders observed in the HBsAG+ set (29%) is intermediate between that of background and vaccine-specific sequences, despite non-specific responders having a substantially larger average cluster size than sequences from either of these classes (89.3), indicating that the method is capturing a subset that is truly enriched with vaccine-specific sequences.

The average abundance of all sequences classified as vaccine specific which are also found in HBsAG+ is similar to the average abundance of all vaccine specific sequences (10.7 in comparison to 10.8). In contrast, in the background and non-specific categories, the average sequence abundance is far higher for those sequences which are also present in the HBsAG+ data set (an increase from 2.62 to 3.45 in background sequences, and 89.3 to 147.1 in vaccine specific sequences). This further suggests that the sequences identified as vaccine specific which are also found in the HBsAG+ data set are truly binding the antigen rather than being selected at random with a size bias.

We next looked at sequence similarity *between* clusters within each class of sequences. Using the Levenshtein distance, we found that sequences classified as vaccine specific were significantly more similar to each other than those of sequences classified as background sequence (*p <* 0.001 based on 1,000 simulations; supplementary material). This is further illustrated in petri-dish plots (Figure 3); here cluster centres were connected by edges if their Levenshtein distance was less than 20% of the sequence length in order to highlight the greater degree of sequence similarity in vaccine specific sequences. Vaccine specific sequences show cliques, and filament structures suggestive of directional selection, while non-responders and particularly background clusters show much less between-cluster similarity.

**Figure 3:**
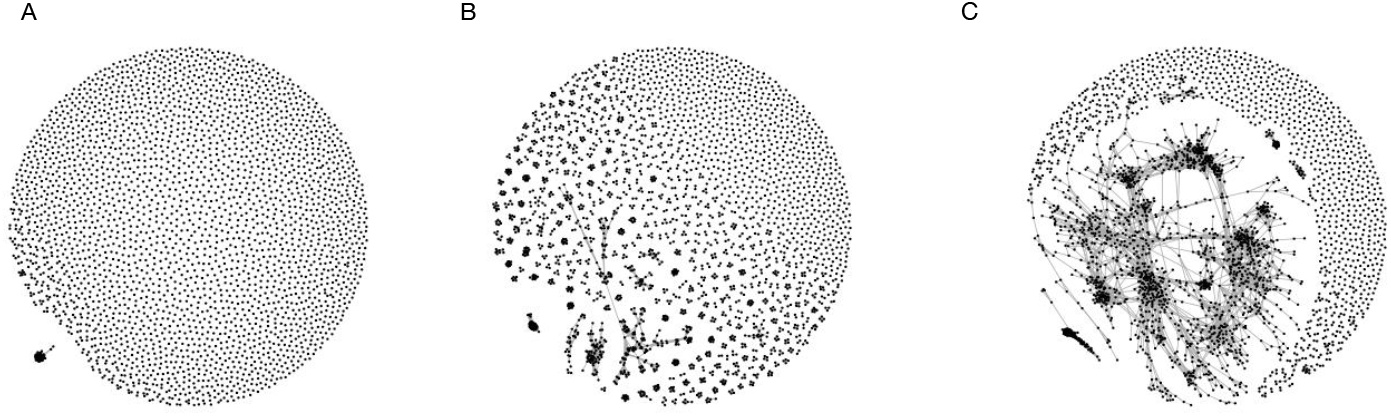
Similarity between sequences classified as background (A), non-specific response (B) and vaccine-specific (C). Each point corresponds to a sequence; sequences are connected if their Levenshtein distance is less than *n/*5 where *n* is the sequence length. All vaccine-specific sequences are shown and a length-matched, random sample of the same number of sequences from the background and non-specific sequences are shown.

### Influenza data set

Fitting the model to the Influenza data set, we again obtain a good QQ plot (see Figure S4) indicating an acceptable model fit, despite considerable differences in the two data sets. Key parameter estimates and an overview of the classification results are given in Tables 3 and 4, and again show that most sequences are classified as belonging to the background population, with only a small fraction classified as responding to any stimuli. However, in this data set, sequences classified as vaccine specific are no more likely to be seen in multiple individuals than those classified as background. Another difference is that the model assigns vanishing weight to the possibility that background sequences are observed at high abundance.

**Table 3:** Fitted parameters to the influenza data set.

| class | $\Gamma_{class}$ | | | $p_{class}$ | | $\omega_{class}$ | |
| --- | --- | --- | --- | --- | --- | --- | --- |
| | bg | ns | vs | bg; ns | vs | bg; $vs_{t=0}$ | ns; $vs_{t>0}$ |
|  | .947 | .001 | .051 | .144 | .144 | 0 | .486 |

**Table 4:** Number of sequences allocated to each category across all samples, the mean total sequence abundance across all samples, and number of sequences also found in the plasmablast data set from each classification.

| Classification | All sequences |  | Plasmablast |
| --- | --- | --- | --- |
|  | Number | Abundance (sd) | Number |
| Background | 27,120 | 1.45 (1.06) | 11 |
| Non-specific | 23 | 5.52 (0.85) | 0 |
| Vaccine-specific | 1,463 | 2.51 (1.54) | 3 |

The sequence abundance and number of individuals sharing sequences over time are illustrated in Figure 4, for each classification. The vaccine specific sequences show a distinct sequence abundance profile, with a sharp increase post-vaccination which reduces over time, whereas the background sequences show little change over time. The average number of individuals sharing a sequence is below one for all categories at all time points, indicating that most sequences are only seen in single individuals and not at multiple time points.

**Figure 4:**
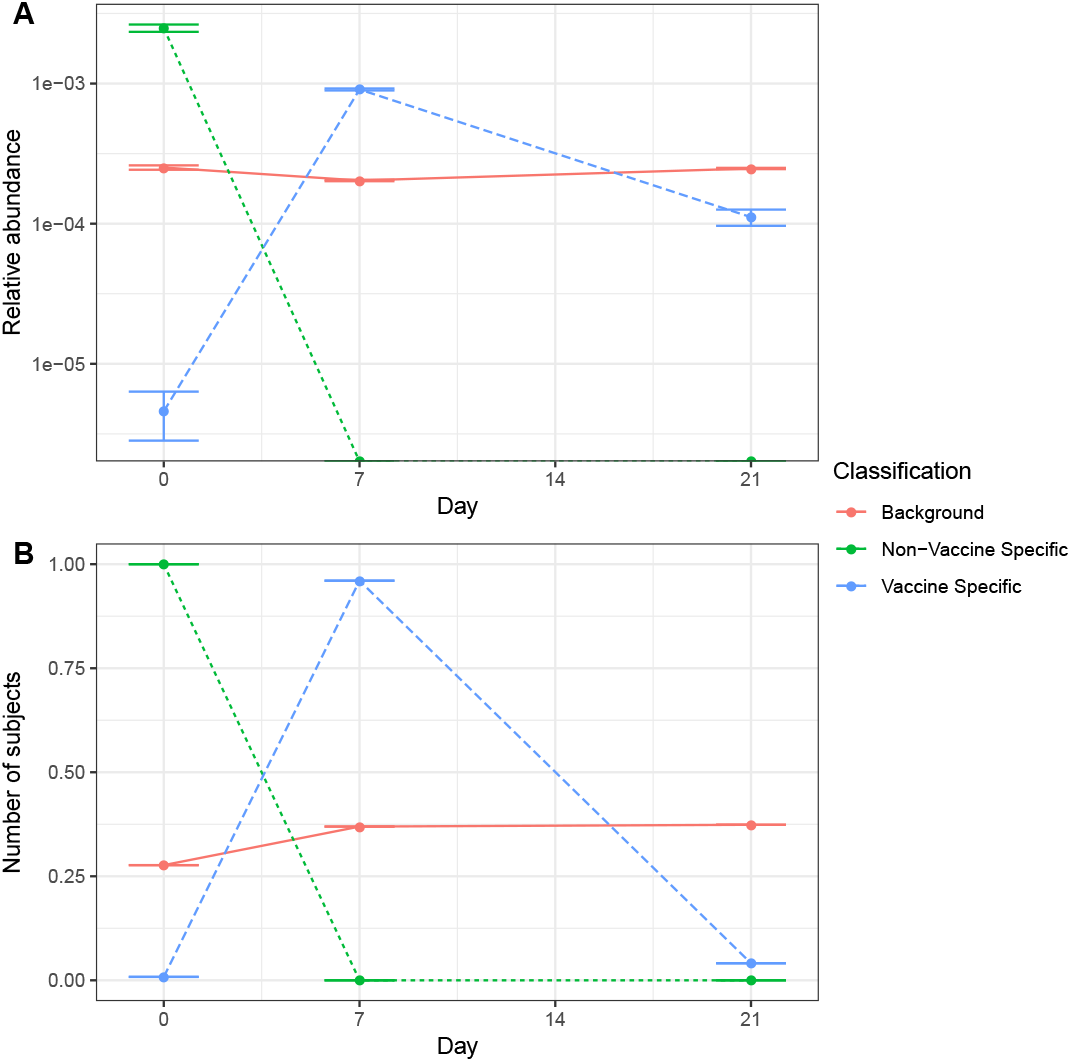
Mean sequence relative abundance at each time point in each classification (A), and the mean number of individuals sharing a sequence over time in each classification (B) for the influenza data set.

The number of sequences allocated to each class and the sequence abundance within each class are shown in Table 4. The majority of sequences are classified as background with a small number being classified as vaccine specific, and only 23 classified as being part of a non-specific response. The sequences classified as vaccine-specific are also typically more abundant.

We then compared the sequences in the influenza data set to those obtained from plasmablasts collected post vaccination. Again, a sequence from the influenza data set was considered to be present in the plasmablast data set if there exists a sequence in the plasmablast data set which would be assigned to the same cluster (Table 2). Of the 436 sequences in the plasmablast data set, 14 are found to be present in the influenza data set, of which 3 would be classified as vaccine specific. These results are considerably less striking as for the hepatitis B data set, although vaccine-specific sequences are still borderline significantly enriched within the monoclonal antibody sequences compared to background sequences (*p* = 0.03, two-tailed Chi-squared test).

The sequences classified as vaccine specific in the influenza data set were also found to be more similar than expected by random chance (*p <* 0.001 based on 1,000 simulations; see Supplementary material). This is illustrated in Figure 5 in which sequences (represented by points) are joined if their Levenshtein distance is less than *n/*3, where *n* is the sequence length. Note that this threshold was chosen to highlight the greater sequence similarity present in vaccine specific sequences and is more stringent than that used for the hepatitis B data set because the viral data consist of amino acid sequences.

**Figure 5:**
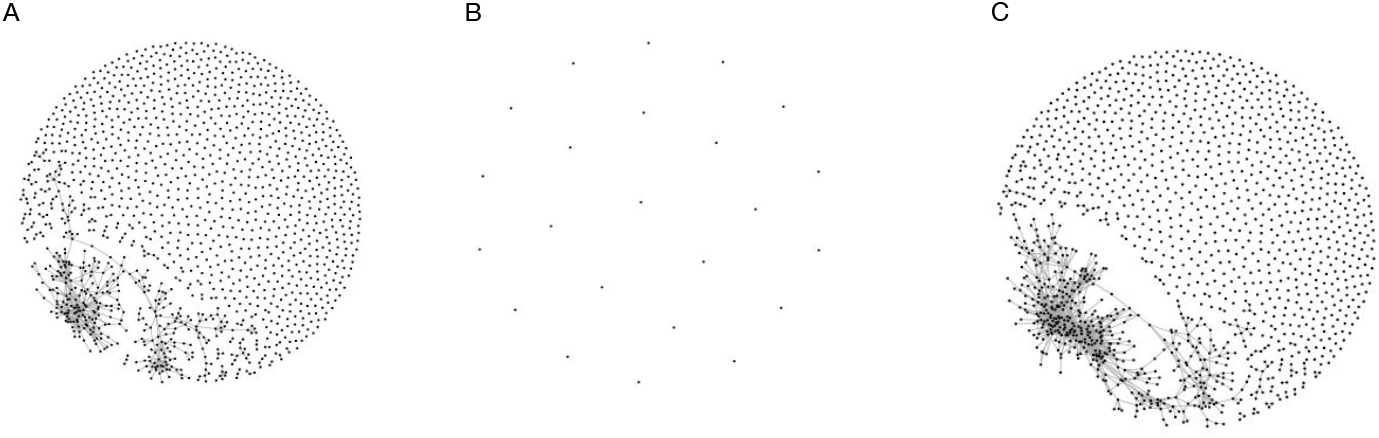
Similarity between sequences classified as background (A), non-specific response (B) and vaccine-specific (C). Each point corresponds to a sequence; sequences are connected if their Levenshtein distance is less than *n/*3 where *n* is the sequence length. All vaccine-specific and non-specific sequences are shown and a random sample from the background which is length and size matched with the vaccine-specific sequences.

## Discussion

The B cell response to vaccination is complex and is typically captured in individuals who are also exposed to multiple other stimuli. Therefore distinguishing B cells responding to the vaccine from the many other B cells responding to other stimuli or not responding at all is challenging. We introduce a model that aims to describe patterns of sequence abundance over time, convergent evolution in different individuals, and the sampling process of B cells, most of which occur at low abundance, from BCR sequences generated pre- and post-vaccination. These patterns are different between B cells that respond to the vaccine stimulus, B cells that respond to a stimulus other than the vaccine, and the bulk of non-responding B cells. By using a mixture model to describe the pattern of sequence abundance for each of these cases separately, we are able to classify sequences as either background, non-specific or vaccine specific.

Vaccine specific sequences are identified with an estimated 69% sensitivity, based on sequences classified as vaccine specific in the hepatitis B data set and their concordance with sequences experimentally identified as vaccine specific in the HBsAG+ data set. The HBsAG+ data set is more likely to contain those sequences present in high abundance in the whole repertoire, due to random chance and a relatively low specificity. This is reflected in the sequences classified as background and as non-specific, in which the average abundance of sequences seen in these categories and in the HBsAG+ data set is higher than the average abundance of all sequences in these categories. However, this over representation of highly abundant sequences is not seen in the cells classified as vaccine specific, suggesting they are indeed binding the vaccine and supporting our estimate of sensitivity.

The influenza data set was compared to the set of sequences from plasmablasts collected post vaccination. However, only 14 of these plasmablast sequences were identified in the influenza set making any estimate of sensitivity from this data set unreliable. Of these plasmablast sequences, 21% were classified as vaccine specific; this is a similar amount to those identified by [10] as in clonally expanded lineages and therefore likely to be responding to the vaccine.

This model incorporates both the signal of sequence abundance as well as sharing between individuals. Considering either of these signals in isolation would not be sufficient to identify vaccine-specific sequences. The sequences we identify as vaccine specific are often highly abundant, but the average sequence abundance is modest, with the non-specific response category containing the most abundant sequences. Of these plasmabalst sequences, 21% were classified as vaccine specific; this is a similar amount to those identified by [10] as in clonally expanded lineages and therefore likely to be responding to the vaccine.

This model incorporates both the signal of sequence abundance as well as sharing between individuals. Considering either of these signals in isolation would not identify as many vaccine-specific sequences with a high sensitivity. Although the sequences we identify as vaccine specific are often highly abundant, their average sequence abundance is modest, with the non-specific response category containging the most abundant sequences. Similarly whilst some sequences identified as vaccine specific were shared between multiple individuals, many were only seen in a single participant. It is only by combining these two signals through the use of a flexible model that we are able to identify the more subtle signatures of vaccine response.

We see evidence for convergent evolution in the hepatitis B data set, with those sequences identified as vaccine specific being much more likely to be seen in multiple individuals. Despite a convergent response to the influenza vaccine being observed by others [10, 17], this pattern is not seen in the influenza data set, in which the probability of a vaccine specific sequence being observed in an individual is similar to that for the background sequences. There are several potential explanations for this. Firstly, in the influenza data set, the signal of sharing among individuals may have been overwhelmed by the abundance signal; many more potentially vaccine specific cells are identified here than in previous studies. Secondly, the influenza data set captures a smaller number of sequences from DNA, whereas the hepatitis B data set captures a larger number of sequences from RNA, so there may be less sharing present in the influenza data set in part due to random chance and in part due to the lack of overrepresentation of highly activated (often plasma cells) B cells. Thirdly, the hepatitis B vaccine was administered as a booster whereas the influenza was a primary inoculation, therefore some optimisation of the vaccine antigen binding is likely to have already occurred after the initial hepatitis B vaccine, increasing the chance that independent individuals converge upon the same optimal antigen binding. Lastly, the complexity of binding epitopes of either of the vaccines is unknown, and the lack of convergent evolution could be explained by a much higher epitope complexity of the influenza vaccine compared to that of the hepatitis B vaccine. This would result in a more diffuse immune response on the BCR repertoire level, making it harder to identify.

In both the hepatitis B and the influenza data sets, it is likely that the sequences show more underlying structure than is accounted for using our clustering approach which only considers highly similar sequences of the same length. The sequences identified as vaccine specific show greater similarity than expected by random chance when utilising the Levenshtein distance, which allows for sequences of different lengths. A possible explanation for this is that there could be a motif shared between sequences of different lengths which could be driving binding specificity. It is possible that by allowing for more complex similarity relationships, larger groups which are more obviously responding to the vaccine may emerge, however current methods are too computationally intensive to allow for complex comparisons of all sequences from all samples.

Here we focus on the signals of sequence abundance and sharing between individuals to identify vaccine specific sequences. The flexibility of the model allows for data sets to be analysed which differed in vaccination strategy, sampling time points, sequencing platforms and nucleic acids targeted. However there are many sequences which are likely incorrectly classified, for instance since random PCR bias can result in large numbers of sequences, if these occur in samples taken at the peak of the vaccine response, they would likely be incorrectly labelled as vaccine specific. Alternatively, vaccination may trigger a non-specific B cell response, B cells involved in this response would have an abundance profile which follows that expected of sequences responding to the vaccine and would therefore likely be misclassified. The inclusion of additional signals, such as hyper-mutation, would improve our model and our estimates of sensitivity.

